# Anti-inflammatory role of GM1 and modulatory effects of gangliosides on microglia functions

**DOI:** 10.1101/2020.03.04.975862

**Authors:** Danny Galleguillos, Qian Wang, Noam Steinberg, Gaurav Shrivastava, Kamaldeep Dhami, Karin Rubinstein, Fabrizio Giuliani, Matthew Churchward, Christopher Power, Kathryn Todd, Simonetta Sipione

## Abstract

Gangliosides are sialic acid-containing glycosphingolipids highly enriched in the brain. Located mainly at the plasma membrane, gangliosides play important roles in signaling and cell-to-cell communication. Lack of gangliosides causes severe early onset neurodegenerative disorders, while more subtle deficits have been reported in Parkinson’s disease and in Huntington’s disease, two misfolded protein diseases with a neuroinflammatory component. On the other hand, administration of ganglioside GM1 provides neuroprotection in both diseases and in several other models of neuronal insult. While most studies have focused on the role of endogenous gangliosides and the effects of exogenously administered GM1 in neurons, their contribution to microglia functions that are affected in neurodegenerative conditions is largely unexplored. Microglia are the immune cells of the brain and play important homeostatic functions in health and disease. In this study, we show that administration of exogenous GM1 exerts a potent anti-inflammatory effect on microglia activated with LPS, IL-1β or upon phagocytosis of latex beads. These effects are partially reproduced by L-t-PDMP, a compound that stimulates the activity of the ganglioside biosynthetic pathway, while inhibition of ganglioside synthesis with GENZ-123346 increases microglial transcriptional response to LPS. We further show that administration of GM1 increases the uptake of apoptotic bodies and latex beads by microglia, as well as microglia migration and chemotaxis in response to ATP. On the contrary, decreasing microglial ganglioside levels results in a partial impairment in both microglial activities. Finally, increasing cellular ganglioside levels results in decreased expression and secretion of microglial brain derived neurotrophic factor (BDNF). Altogether, our data suggest that gangliosides are important modulators of microglia functions that are crucial to healthy brain homeostasis, and reveal that administration of ganglioside GM1 exerts an important anti-inflammatory activity that could be exploited therapeutically.

## INTRODUCTION

Microglia, the innate immune cells of the central nervous system (CNS), play important homeostatic roles in health and disease. Initially identified as the orchestrators of the CNS immune response in the context of brain injury or disease, it is now clear that during development and in the healthy adult CNS microglia exert modulatory and housekeeping functions [1, 2] that span from synapse remodeling and maturation [3, 4] to secretion of neurotrophic factors [5, 6] and regulation of the pool of neuronal precursors [7]. With ageing, homeostatic and neurotrophic activities of microglia decline or become progressively impaired [8], promoting the onset and/or progression of neurodegenerative conditions [9-11]. A maladaptive increase in microglia inflammatory responses, often associated with a decrease in reparative and neurotrophic functions, occurs in common neurodegenerative diseases, including Alzheimer’s disease (AD) [12], Huntington’s disease (HD) [13, 14] and Parkinson’s disease (PD) [15, 16]. Therefore, treatments that curtail detrimental microglia activities while boosting neuroprotective ones in pathological conditions could be highly beneficial.

Glycans play a major role in the regulation of immune cell functions [17]. The glycome of the CNS is unique in that it is predominantly composed of glycolipids - as opposed to glycoproteins in peripheral systems [18] - with gangliosides being the most abundant species [19]. Gangliosides are sialic acid-containing glycosphingolipids enriched at the plasma membrane and play key modulatory roles in signaling and cell-to-cell communication [20]. Their importance for brain health is highlighted by the fact that loss of function mutations that affect their synthesis causes neurodegeneration in humans [20-26] and mice [27]. A decrease in ganglioside levels and changes in the relative abundance of specific gangliosides also occur in ageing [28-31] and in common neurodegenerative conditions, including HD [32, 33], PD [34] and AD [35, 36]. On the other hand, therapeutic administration of one of the most abundant brain gangliosides, GM1, was shown to provide neuroprotection in models of neuronal injury and neurodegeneration [37-40] and in genetic models of HD [41, 42].

The mechanisms underlying the neuroprotective effects of endogenously synthesized as well as therapeutically administered gangliosides are only partially understood, with past studies having focused mainly on their effects in neurons. Their roles in other brain cells remain largely unexplored or controversial. The few *in vivo* studies that have investigated the effects of lack of gangliosides or exogenous ganglioside administration on microglia activation and neuroinflammation are difficult to interpret, due to concomitant confounding effects in neuronal cells [43-45]. It is also unknown whether changes in endogenous ganglioside levels, as observed in ageing and disease, or administration of exogenous gangliosides for neuroprotection might contribute to skew microglia towards inflammatory or reparative/protective phenotypes, respectively. Consequently, direct investigation of the effects of gangliosides on isolated microglia is crucial to determine whether they play a modulatory role on microglia functions that might contribute to brain health and disease, and to elucidate the neuroprotective effects of therapeutically administered gangliosides.

In this study we used two complementary experimental paradigms to address these questions: 1) administration of exogenous GM1 to determine the potential effects of therapeutically administered ganglioside on microglia and neuroinflammation; 2) modulation of microglial ganglioside levels through the use of a pharmacological activator and an inhibitor of the ganglioside biosynthetic pathway, the latter to mimic the partial decrease in ganglioside levels that has been observed in some neurodegenerative conditions [32-34]. We demonstrate that administration of exogenous ganglioside GM1 triggers a deactivating mechanism in murine and human microglia that curtails the inflammatory response induced by various stimuli, including LPS, IL-1β and engulfment of latex beads. Furthermore, modulation of endogenous ganglioside synthesis significantly affects microglia response to LPS, microglia ability to clear apoptotic bodies and to migrate in response to chemotactic stimuli. Altogether, our results suggest that gangliosides play an important role in microglia physiology and that modulation of their levels in microglia can offer new therapeutic avenues in neurodegenerative and neuroinflammatory diseases.

## METHODS

### Animal and human studies

All procedures on animals were approved by the University of Alberta Animal Care and Use Committee and were in accordance with the guidelines of the Canadian Council on Animal Care. Human primary microglia cultures were prepared from fetal tissue obtained from15-20 week electively terminated healthy pregnancies with written informed consent of the donors (Pro000027660), as approved by the University of Alberta Human Research Ethics Board (Biomedical).

### Chemicals and reagents

Ganglioside GM1 (purified from porcine brain) was obtained from TRB Chemedica Inc. (Switzerland) and resuspended in cell culture grade water. Lipopolysaccharide (LPS, serotype O55:B5, gamma-irradiated) was purchased from Sigma (Sigma L6529), recombinant mouse IL-1β was purchased from Cedarlane (Cedarlane CLCYT273), L-*threo*-1-phenyl-2-decanoylamino-3-morpholino-1-propanol•HCl (L-*t*-PDMP) was from Matreya LLC (Matreya #1749) and was dissolved in ethanol, and N-[(1R,2R)-1-(2,3-Dihydrobenzo[b][1,4]dioxin-6-yl)-1-hydroxy-3- (pyrrolidin-1-yl)propan 2-yl] nonanamide (GENZ-123346) was obtained from Toronto Research Chemicals (TRC G363450) and solubilized in DMSO. All other reagents were from Sigma, unless otherwise indicated.

### Primary cultures and cell lines

Primary mixed glial cultures from P0.5-P1.5 mice (C57BL/6J or FVB/NJ strain) and rats (Sprague Dawley) were prepared following the method described by [46]. In brief, cerebral cortices were enzymatically and mechanically dissociated and cells were seeded in DMEM/F12 medium supplemented with 10% FBS, 100 units/ml penicillin, 100µg/ml streptomycin, 1 mM sodium pyruvate and 50 µM β-mercaptoethanol. Growth medium was replaced every 4 days. On day 14, the medium was removed, and cultures were trypsinized to remove the monolayer of astrocytes leaving adherent microglia attached to the bottom of the culture dish. Isolated microglia was left to rest for 24 hours in DMEM/F12 without FBS (supplemented with 1 mM Sodium Pyruvate and 50 µM β-mercaptoethanol) before any experimental treatment. Alternatively, microglia on top of mixed glial cultures were harvested by incubation with 15 mM lidocaine in growth medium in a shaking incubator for 15 min at 200 rpm [47]. Human fetal microglia were prepared as previously reported [48], from human fetal tissue obtained from 15-20 wk electively terminated healthy pregnancies with the written informed consent (Pro000027660), as approved by the University of Alberta Human Research Ethics Board (Biomedical). Briefly, fetal brain tissue was dissected, meninges were removed, and a single cell suspension was prepared by enzymatic digestion followed by passage through a 70-μm cell strainer. Cells were plated in T-75 flasks and maintained in MEM supplemented with 10 % FBS, 2 mM L-glutamine, 1 mM sodium pyruvate, MEM nonessential amino acids, 0.1 % dextrose, 100 U/ml Penicillin, 100 μg/ml streptomycin, 0.5 μg/ml amphotericin B, and 20 μg/ml gentamicin. Mixed cultures were maintained for 2 weeks. Weakly adhering microglia were recovered by gently rocking the mixed cultures for 20 min, followed by cell decanting, washing and plating onto 96 well plates (50,000 cells/well). Isolated microglia were allowed to rest for 3 days prior to performing experiments. BV-2 cells cells [49] (kindly donated by Dr. Jack Jhamandas, University of Alberta) were grown in RPMI-1640 supplemented with 10% FBS, 1 mM Sodium Pyruvate, 2 mM L-glutamine and 50 µM β-mercaptoethanol. Neuro-2a cells (N2a, kindly donated by Dr. Satyabrata Kar, University of Alberta) were grown in DMEM: Opti-MEM 1:1 containing 10% FBS supplemented with 1 mM Sodium Pyruvate and 2 mM L-glutamine. All cells were maintained at 37°C in 5% CO_2_.

### Cell treatments

GM1 was applied to primary microglia concomitantly with LPS or after washing off LPS from the cells. For the first protocol, after isolation and 24 hours of resting in medium without FBS, the medium was removed and GM1 50µM dissolved in DMEM/F12 supplemented with 1 mM Sodium Pyruvate and 50 µM β-mercaptoethanol was added to the cells. After 1 hours, LPS was added directly to the medium at a final concentration of 100ng/ml. Treatments with IL-1β were performed in a similar manner, replacing LPS with IL-1β (5ng/ml). For the second set of experiments, the cells were left to rest after isolation as described and after 24 hours the medium was removed and LPS (100ng/ml) dissolved in DMEM/F12 supplemented with 1 mM Sodium Pyruvate and 50 µM β-mercaptoethanol was added to the cells. After 3 hours of stimulation with LPS, the medium was removed and the cells were washed once with HBSS^++^ containing 0.1% essentially fatty acid free BSA. After that, cells were washed one more time with HBSS^++^ and twice with DMEM/F12. Immediately after, GM1 50µM dissolved in DMEM/F12 supplemented with 1 mM Sodium Pyruvate and 50 µM β-mercaptoethanol was added to the cells.

### Immunoblotting

BV-2 cells or primary microglia were lysed in ice-cold 20 mM Tris, pH 7.4, containing 1% Igepal CA-630, 1 mM EDTA, 1 mM EGTA, 1X cOmplete protease inhibitor and PhosStop phosphatase inhibitor cocktails (Roche) and 50 µM MG-132. For immunoblotting analysis, 20 μg of proteins were separated by 10% SDS-PAGE and transferred to Immobilon-FL membranes (Millipore). For dot-blot assays, 2 μg of proteins were spotted onto nitrocellulose membranes (Bio-Rad) using a dot-blotting apparatus (Bio-Rad) according to manufacturer’s instruction. After blocking in Odyssey blocking buffer (LI-COR) for 1h, the membranes were incubated overnight at 4°C with the following primary antibody: rabbit anti-IKKα (Cell Signaling 2682; 1:1,000), rabbit anti-phospho-IKKα/β (Ser180/Ser181) (Cell Signaling 2681; 1:1,000), rabbit anti-p38 MAPK (Cell Signaling 9212; 1:1,000), rabbit anti-phospho-p38 MAPK (Thr180/Tyr182) (Cell Signaling 9211; 1:1,000), mouse anti-α-Tubulin (Sigma T5168; 1:5,000), rabbit anti-GM1 (Calbiochem 345757: 1:1,000), mouse anti-GD1a (Millipore MAB5606Z; 1:1,000), mouse anti-GD1b (DSHB GD1b-1 1:200), mouse anti-GT1b (Millipore MAB5608; 1:500) or cholera toxin subunit B-Alexa647 (Invitrogen C34778, 1µg/ml), followed by appropriate IRDye secondary antibody (LI-COR, 1:10,000) for 1h at room temperature. Infrared signals were acquired and quantified using an Odyssey Imaging System (LI-COR) instrument.

### RNA extraction and qPCR analysis

Primary microglia or BV-2 cells were collected in RLT buffer (QIAGEN). RNA was isolated using RNEasy Micro or Mini Kit, respectively. cDNA was synthetized from 200-500 ng of RNA reverse transcribed using Oligo dT primers and SuperScript II (Invitrogen). qPCR was carried out using PowerUp SYBR Green Master Mix (Applied Biosystems) in a StepOne Plus instrument (Applied Biosystems). Gene expression was normalized over the geometric mean of the expression of 3 reference genes: Atp5b, Cyclophilin A and Rplp0 (Normalization Index), according to [50].

### ELISA

Cytokines released by primary microglia in the culture medium were quantified by ELISA using the following commercial kits according to manufacturer’s instruction: Mouse TNF alpha Uncoated ELISA kit (88-7324-22, Invitrogen), Rat IL-6 DuoSet ELISA (DY506) and Rat IL-1β DuoSet ELISA (DY501) (R&D Systems), and Human IL-1β DuoSet ELISA (DY201, R&D Systems). BDNF and IGF-1 in cell medium and cell lysates of mouse primary microglia were measured using Mature BDNF Rapid ELISA kit (Biosensis, BEK-2211-1P) and Mouse/Rat IGF-1 Quantikine ELISA kit (R&D Systems, MG100), respectively. Cytokines and growth factors levels were normalized to total protein content in the corresponding cell lysates.

### Chemotaxis assay

Chemotaxis of primary microglia towards ATP was assessed using a transwell migration assay (REF?). Briefly, FVB/NJ primary microglia harvested from mixed glia cultures by the shaking method were seed onto Millicell cell culture inserts with 8-μm pore filter (Millipore), and treated with 50 μM GM1 for 4h or with 10 μM L-*t*-PDMP or 5 μM GENZ-123346 for 3 days. The bottom wells were then filled with medium containing 100 μM ATP (Sigma), and microglia were allowed to migrate from the top side to the bottom side of the filter for 2h at 37°C. Non-migrating cells on top of the filter were wiped off using a Q-tip, while migrated microglia on the bottom side were stained with 0.5µg/ml DAPI for 5 min. Images of the filters were collected using a Zeiss AxioVision fluorescence microscope and cell nuclei were counted using ImageJ software.

### *In vitro* scratch assay

*In vitro* scratch assay for performed as described [51]. Briefly, primary microglia or BV-2 cells were cultured on poly-l-lysine-coated 48-well plates at 37°C overnight. After treatment with GM1 (50 μM) for 4h, the cell monolayer was gently scratched with a sterile 200 µl pipet tip and detached cells were wash away with medium. Cells were incubated at 37°C to allow migration into the cell-free region. The scratch area was photographed using Zeiss AxioVision microscope and migrating cells into the area were quantified using ImageJ software.

### Phagocytosis assay

To prepare apoptotic bodies, neuronal N2a cells were cultured overnight with 0.5 μM staurosporine to induce apoptosis. Apoptotic bodies and cell debris were collected by centrifugation at 2,000 x g for 10 min and then labelled with 1µg/ml pHrodo Red (Invitrogen P36600) in freshly prepared 100 mM bicarbonate buffer (pH 8.5), for 1h at rt, protected from light. Microglia were pre-treated with 50 μM GM1 for 4h or with 10 μM L-*t*-PDMP or 5 μM GENZ-123346 for 3 days, then washed with HBSS containing 0.1%BSA and incubated at 37°c for 24 h with equal amounts of pHrodo-labeled apoptotic bodies in HEPES-buffered and phenol red-free DMEM/F12 medium. Fluorescence emission of pHrodo in the acidic phagolysosome compartment of microglia cells upon phagocytosis was quantitated using a SpectraMax microplate reader (Molecular Devices). To evaluate phagocytic activity towards latex beads, the Phagocytosis Assay Kit (IgG FITC) (Cayman Chemical 500290) was used according to the manufacturer’s instruction. Briefly, primary microglia or BV-2 cells were incubated with 1µm FITC-beads for 2h at 37°C, unbound beads were washed away with cold PBS and trypan-blue was added for 2 min at room temperature to quench the fluorescence of any remaining bead on the surface of microglia. Phagocytosis of latex beads by primary microglia co-labelled with goat anti-Iba-1 antibodies (Abcam ab107159) was performed in the High Content Analysis Core of the Faculty of Medicine & Dentistry (University of Alberta) using an Operetta high content imaging system (PerkinElmer). Phagocytic activity was defined as the mean integrated fluorescence (MIF) intensity of cells that had taken up at least 1 bead. Phagocytosis of latex beads by BV-2 cells was quantified by flow cytometry in the Flow Cytometry Core Facility of the Faculty of Medicine & Dentistry at the University of Alberta, and analyzed with FlowJo software.

### Statistical analysis

Two tailed *t*-test analysis, One-way ANOVA corrected for multiple comparisons (Sidak’s post-test) or two-way ANOVA corrected for multiple comparisons (Tukey’s post-test) were used where appropriate using GraphPad Prism 7.

## RESULTS

### Administration of exogenous GM1 decreases microglia activation following pro-inflammatory stimulation with LPS

To determine the specific effects of GM1 on microglia in inflammatory conditions, BV2 microglia cells were pre-incubated with GM1 (50 µM) for 2 h and then stimulated with LPS (E. coli serotype O55:B5, 100ng/ml) to stimulate the Toll-like receptor 4 (TLR4), a major pattern recognition receptor that is also activated by endogenous danger-associated molecular patterns released in stress and neurodegenerative conditions [52-54]. In untreated cells, LPS stimulation induced activation of the NFkB and the MAPK pathways, as expected and as shown by phosphorylation of IKK and p38 MAPK, respectively, within 30 minutes of LPS administration. In cells pre-treated with GM1, this response was significantly attenuated (Figure 1A) and correlated with a dramatic decrease in the downstream expression of NFkB pro-inflammatory target genes, including TNF and IL-1β (Figure 1B). We confirmed these results in primary cultures of rat and mouse microglia, where administration of GM1 prior to a challenge with LPS blocked the release of the pro-inflammatory cytokines IL-6 and IL-1β in the cell culture medium and the synthesis of NO (Figure 1C), as well as transcription of IL-1β, TNF and IκBα (Figure 1D). The effects of GM1 on microglia activation was not due to decreased *surface* expression of the LPS receptor (Toll-like receptor 4, TLR4), although cells treated with GM1 for 24h, but not 4h, did show a decrease in *total* cellular TLR4 compared to untreated cells (Supplementary Figures 1A and 1B), nor to changes in cell viability (Supplementary Figure 2). Furthermore, GM1 did not affect basal expression of pro-inflammatory cytokines.

**Figure 1.**
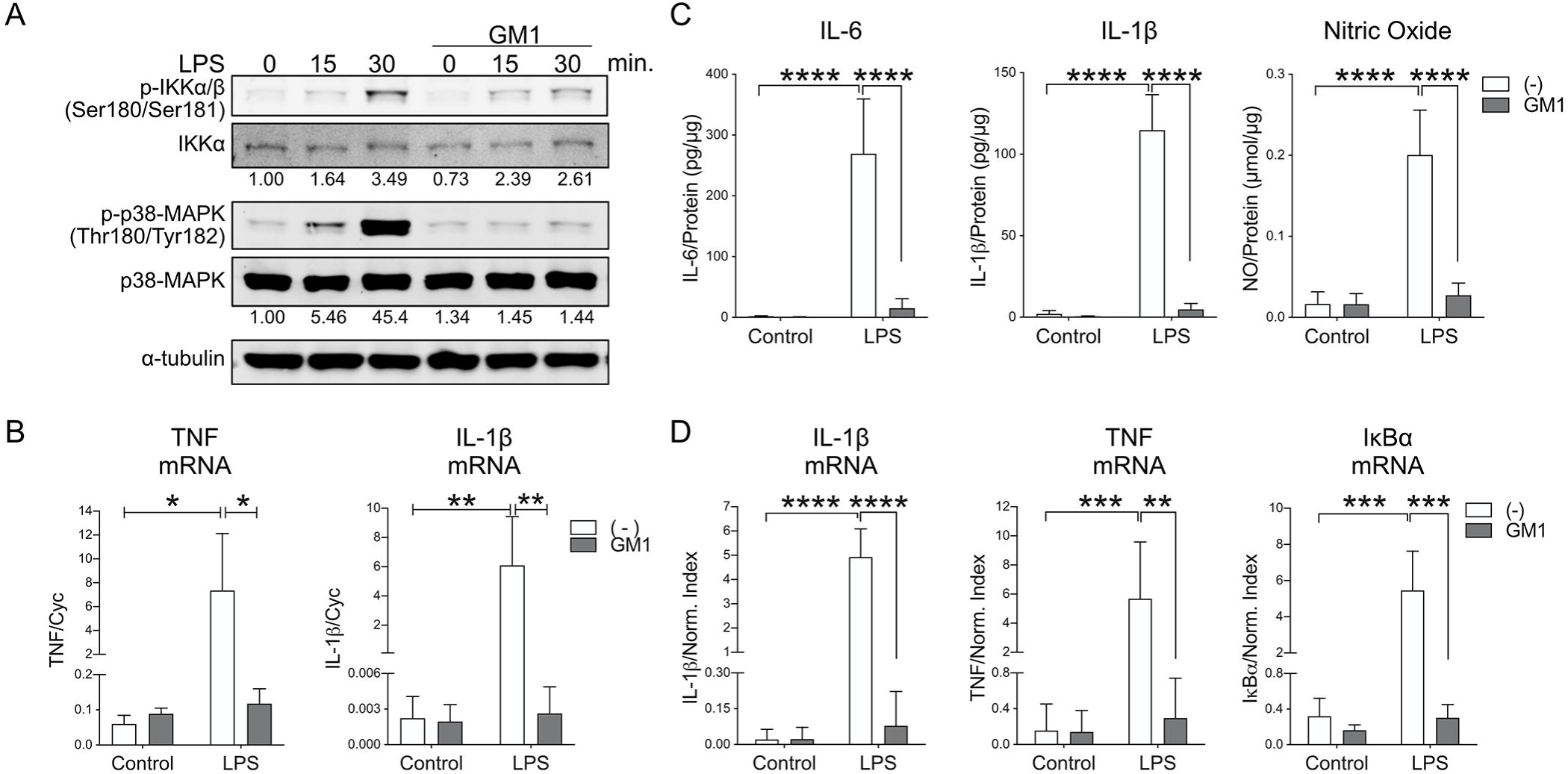
Microglia incubation with ganglioside GM1 prevents pro-inflammatory activation by LPS. **A)** BV2 microglial cells were pre-incubated with GM1 (50µM) or vehicle for 2h prior to stimulation with LPS (100 ng/ml). Representative immunoblot showing decreased levels of phospho-IKKα/β and phospho-p38 MAPK following stimulation with LPS in cells pre-treated with GM1. The numbers under the immunoblots show fold-change over unstimulated controls, normalized over total protein. **B)** Upregulation of TNF and IL-1β mRNA expression in response to LPS stimulation (100 ng/ml for 6h) is blunted in BV-2 cell pre-incubated for 1h with GM1 (N=3). **C)** Pre-incubation with GM1(50 µM; 1 hours) prevents IL-6, IL-1β and nitric oxide (NO) release in the medium by rat microglia stimulated with LPS (100 ng/ml, 24 hours) (N=3). **D)** Mouse primary microglia were treated as in C) and expression of IL-1β, TNF and IκBα mRNA was measured (N=4-5). Bars show mean values ± STDEV. Two-way ANOVA with Tukey’s multiple comparisons test; \**p*<0.05, \*\**p*<0.01, \*\*\**p*<0.001, \*\*\*\**p*<0.0001.

Next, we set up to determine whether GM1 would dampen microglia inflammatory responses when administered to cells *after* stimulation with LPS. Under these conditions, GM1 still decreased TNF and IL-1β gene expression (Figure 2A) as well as the amount of TNF cytokine released into the culture medium (Figure 2B), although to a lesser extent than when administered prior to LPS stimulation. Importantly, the effects of GM1 were not limited to rodent microglia, but extended to human fetal microglia, where secretion of IL-1β upon LPS stimulation was greatly decreased by subsequent treatment with GM1 (Figure 2C).

**Figure 2.**
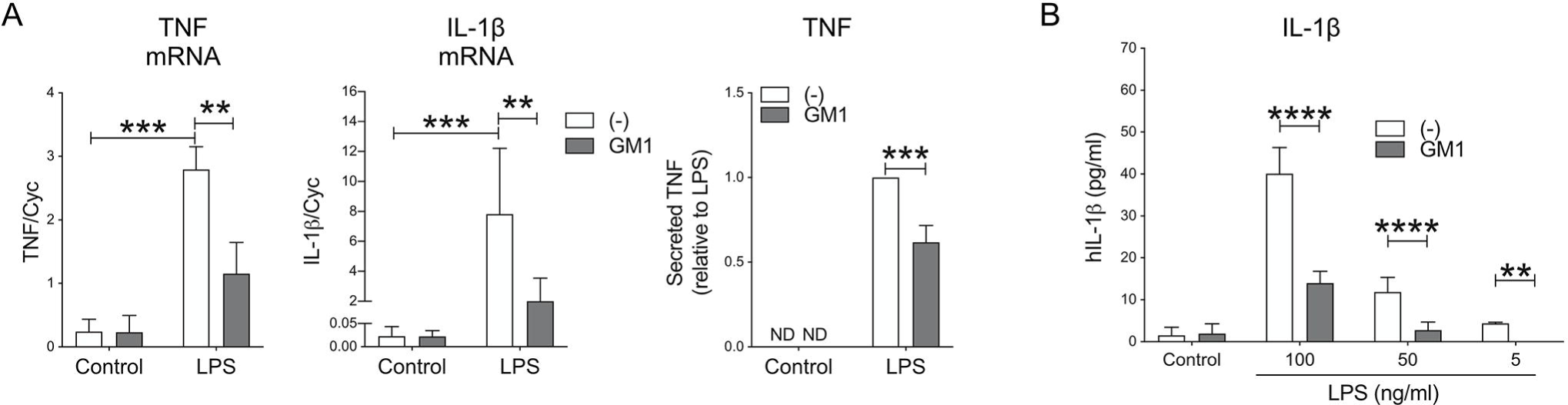
Administration of GM1 after microglia stimulation with LPS curtails pro-inflammatory microglia activation. **A)** Mouse primary microglia were stimulated for 3h with LPS (100 ng/ml), after which LPS was washed out and cells were further incubated with GM1 (50 µM) for 8 h. Expression of TNF and IL-1β mRNA and TNF secreted in the medium were significantly decreased in GM1-treated cells (N=3-5). **B)** Human fetal microglia were treated as in A). GM1 treatment decreased IL-1β secretion into the medium (N=3). Bars show mean values ± STDEV. Two-way ANOVA with Tukey’s multiple comparisons test. \*\**p*<0.01, \*\*\**p*<0.001, \*\*\*\**p*<0.0001.

To determine the effects of GM1 administration on microglia activation triggered by other pro-inflammatory stimuli, we exposed mouse microglia to IL-1β to mimic a physiological response to a milder stimulus relevant to neurodegenerative and neuroinflammatory conditions [55, 56]. Administration of GM1 blocked the NFkB transcriptional response to IL-1β stimulation, as measured by expression of IL-1β (Figure 3A). Furthermore, GM1 decreased the expression of pro-inflammatory genes following exposure of BV2 cells to latex beads, another stimulus known to activate inflammatory responses in microglia [57], without affecting the amounts of phagocytosed beads (Figure 3B). Altogether, these data suggest that administration of GM1 dampens microglial inflammatory responses to different stimuli and can even curtail microglia activation after exposure to inflammatory conditions.

**Figure 3.**
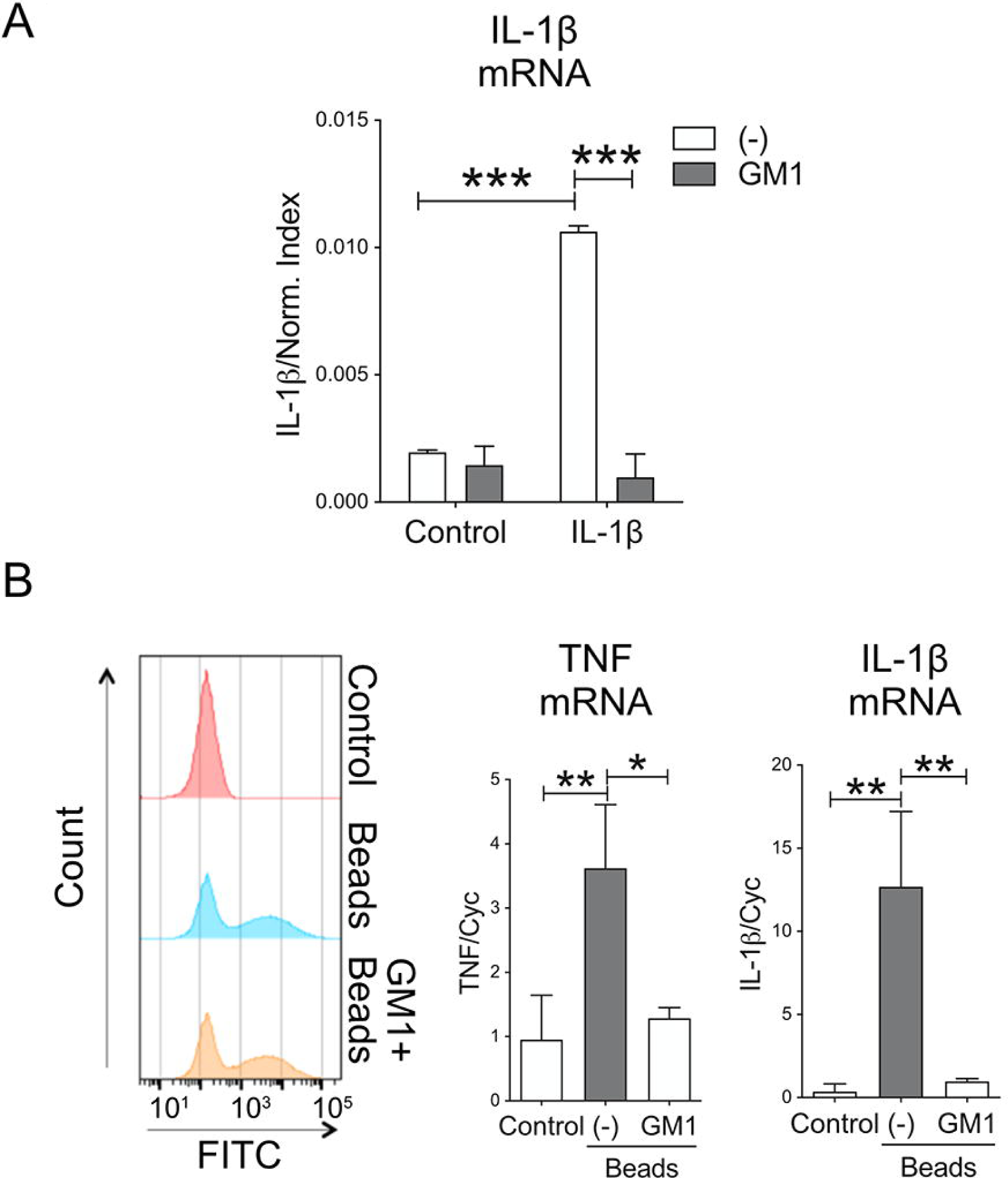
Pre-incubation of microgliawith GM1 decreases pro-inflammatory activation triggered by IL-1β and by phagocytosis of polystyrene beads. **A)** Mouse primary microglia were pre-incubated with GM1 (50 µM; 1 hours) and then stimulated with IL-1β (5 ng/ml) for 24h. GM1 blocked upregulation of IL-1β mRNA expression. **B)** Phagocytosis of latex beads by BV-2 cells pre-treated with GM1 for 2h. Representative histograms of beads uptake are shown. GM1 pre-treatment did not affect uptake Graphs show mRNA expression for IL-1β and TNF upon bead phagocytosis. Data are mean values ± STDEV of 3 independent experiments. Two-way ANOVA with Tukey’s multiple comparisons test was used in A). One-way ANOVA with Sidak’s multiple comparisons test was used in B) Data are mean values ± STDEV from 3 independent experiments. \**p*<0.05, \*\**p*<0.01, \*\*\**p*<0.001.

### Endogenous ganglioside levels modulate the response of microglia to pro-inflammatory stimulation

To determine whether *endogenous* ganglioside levels can affect the response of microglia to pro-inflammatory stimuli, we used the compound L-*t*-PDMP to stimulate the activity of microglial UDP-glucose ceramide glucosyltransferase (UGCG) [58] (Fig. 4A) and increase cellular ganglioside levels. In BV2 cells, treatment with 5 and 15 µM L-*t*-PDMP resulted in a modest but significant increase in the levels of gangliosides GM1 and GT1b (Figure 4B), two of the four most abundant gangliosides in the brain. These changes in ganglioside levels were accompanied by decreased expression of total cellular TLR-4 (Figure 4C), although surface expression was similar between L-*t*-PDMP-treated and untreated cells (Supplementary Figure 1B), and decreased microglia activation upon exposure to LPS (100 ng/ml), as shown by decreased phosphorylation of IKK and p38-MAPK (Figure 4D). Concomitantly, we observed a significant reduction in LPS-induced expression of IL-1β and TNF mRNA (Figure 4E). Similar results were obtained in mouse primary microglia, where stimulation of ganglioside synthesis with L-*t*-PDMP resulted in significantly increased levels of GM1, GD1a and GT1b (Figure 4F) and decreased levels of IL-1β and TNF mRNA at low LPS concentrations (0.01 and 0.1 ng/ml), although these effects were lost at higher LPS concentrations (100 ng/ml) (Figures 4G and 4H). TNF and IL-1β released in the medium were undetectable at the lowest LPS concentrations used, while upon microglia stimulation with 100 ng/ml LPS, the amount of TNF secreted into the medium was similar between groups (Figure 4H), in agreement with gene expression data. Overall, our data suggest that increasing endogenous levels of microglial gangliosides attenuates microglia response to an inflammatory stimulus.

**Figure 4.**
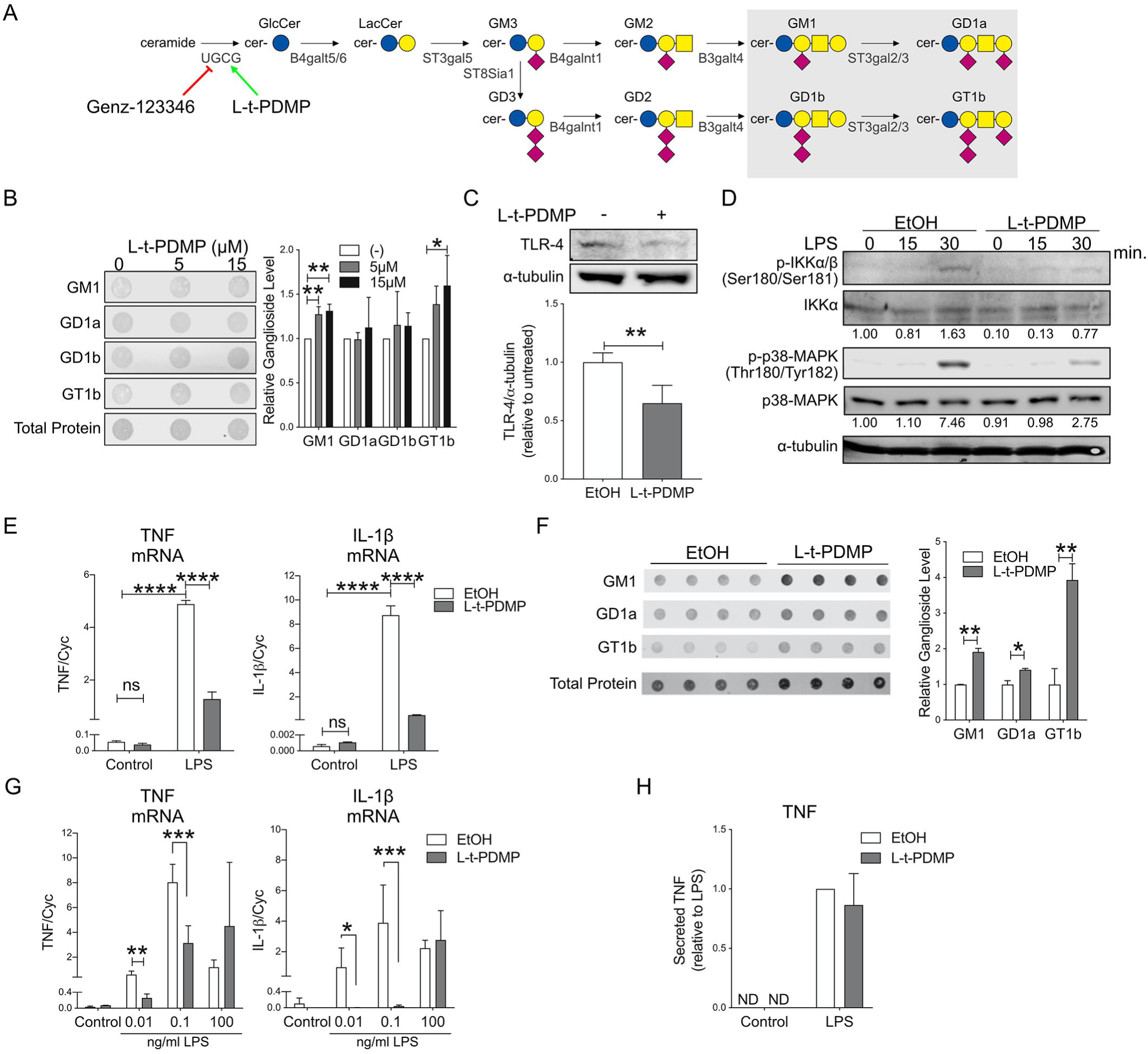
Stimulation of the ganglioside biosynthetic pathway with L-t-PDMP decreases pro-inflammatory microglia activation. **A)** Simplified scheme of the ganglioside biosynthetic pathway and relative enzymes. L-t-PDMP and Genz-123346 are an activator and an inhibitor, respectively of UDP-glucose ceramide glucosyltransferase (UGCG). The shaded grey area highlights the major brain gangliosides. Blue circle: glucose; yellow circle: galactose, yellow Glucose: blue circle, Galactose: yellow circle, N-Acetylgalactosamine: yellow square, N-Acetylneuraminic acid: purple diamond. **B)** BV2 cells were incubated for 72 h with the indicated concentrations of L-t-PDMP to increase ganglioside synthesis. A representative dot blot and quantification of cellular ganglioside levels before and after treatment are shown (N=3). **C)** Representative immunoblot and densitometric analysis of TLR-4 expression showing decreased levels of the receptor in BV2 cells treated with L-t-PDMP (N=6). D) Representative immunoblot showing phospho-IKK and phospho-p38 MAPK levels in BV2 cells stimulated with LPS (100 ng/ml) for the indicated time, after cell treatment with L-t-PDMP (15µM, 72h). The numbers under the blots show fold-change over untreated control, after normalization for total IKK or p38-MAPK levels. The experiment was repeated twice with similar results. **E)** Expression of TNF and IL-1β mRNA in BV-2 cell treated as indicated above (N=3). **F)** Dot-blot analysis of ganglioside levels in murine primary microglia incubated with L-t-PDMP (10µM for 72h). **G)** Expression of IL-1β and TNF mRNA in primary microglia treated for 72 hours with 10µM L-t-PDMP and stimulated with the indicated concentrations of LPS for 6 h (N=3). **H)** TNF secretion by murine microglia after treatment with L-t-PDMP and stimulation with the indicated concentrations of LPS (N=3). One-way ANOVA with Sidak’s multiple comparisons test was used in B); two-tailed *t*-test was used in C and F); two-way ANOVA with Tukey’s multiple comparisons test was used in E, G and H. \**p*<0.05, \*\**p*<0.01, \*\*\**p*<0.001, \*\*\*\**p*<0.0001.

Various studies have shown that brain ganglioside levels are decreased in some common neurodegenerative conditions, including PD and HD [20-26, 32, 34]. To explore the potential impact of these changes on microglia biology and determine whether they could contribute to the development of a neuroinflammatory state in pathological conditions, we used the compound GENZ-123346 [59] to specifically *inhibit* the activity of UGCG in BV2 cells and primary microglia and reduce cellular ganglioside levels. In BV2 cells, treatment with GENZ-123346 resulted in >50% decrease in GM1 and GD1a levels, although, unexpectedly, the amount of the complex ganglioside GT1b increased slightly (Figure 5A). These changes were accompanied by increased phosphorylation of IKK and p38-MAPK upon cell stimulation with LPS (Figure 5C) - despite expression of total and surface TLR-4 expression was not significantly affected by the treatment (Figures 5B and Supplementary Figure 1B) - and in a modest but significant increase in the expression pro-inflammatory cytokines compared to untreated cells (DMSO, Figure 5D). Similar to BV2 cells, treatment of primary microglia with GENZ-123346, resulted in decreased levels of gangliosides GM1 and GD1a (Figure 5E), and decreased transcription of TNF after microglia stimulation with LPS (0.01 ng/ml for 6h), although no effects were observed on IL-1β transcription (Figure 5F). Surprisingly, the increase observed for TNF mRNA was not accompanied by a corresponding increase in the amount of TNF secreted in the medium, which was slightly decreased in cells treated with GENZ-123346 (Figure 5G).

**Figure 5.**
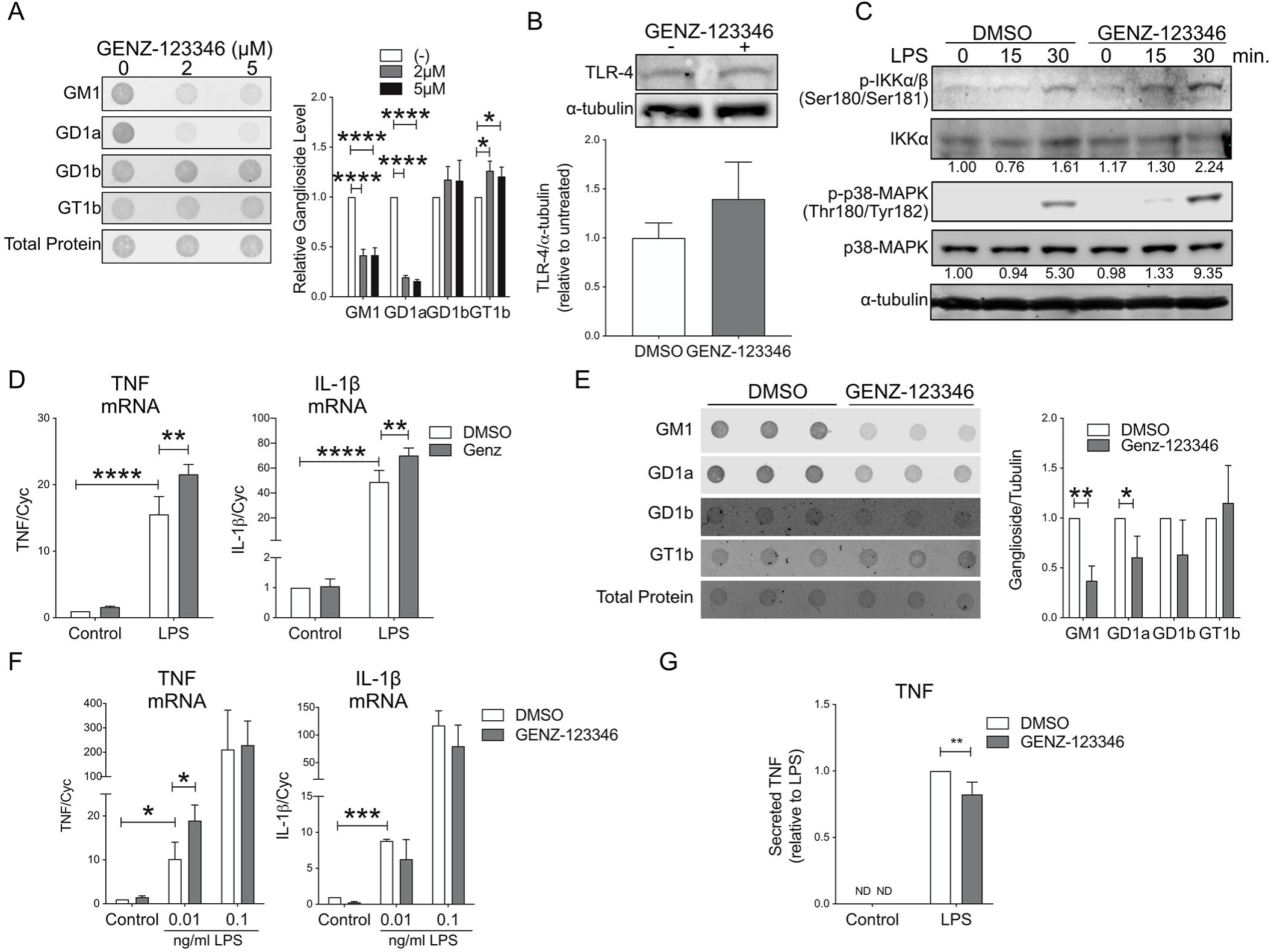
Inhibition of the ganglioside biosynthetic pathway enhances microglial response to LPS. **A)** Dot blot and relative quantification of cellular ganglioside levels in BV2 cells treated with the indicated concentrations of GENZ-123346 for 72 hours (N=3). **B)** Representative immunoblot and quantification of TLR-4 expression in BV-2 cells after 72 h incubation with GENZ-123346 (5µM) (N=5). **C)** Representative immunoblot showing increased phosphorylation of IKK and p38 MAPK in BV-2 cells treated with GENZ-123346 and stimulated with LPS (100 ng/ml) for the indicated time. The numbers under the blots indicate fold-change of p-IKK and p-p38-MAPK compared to untreated control, after normalization over total IKK or p38-MAPK protein levels. **D)**. Expression of IL-1β and TNF mRNA in BV-2 cell stimulated with LPS (100 ng/ml) after 72 h incubation with GENZ-123346 (5μM) (N=3). **E)** Dot blot and quantification of ganglioside levels in primary mouse microglia incubated with 10µM GENZ-123346 for 72h (N=3). **F)** Expression of IL-1β and TNF mRNA in mouse microglia treated with GENZ-123346 as in E), and after stimulation with LPS (0.01 ng/ml for 6h) (N=3). **G)** TNF secretion by control and GENZ-123346-treated cells upon stimulation with LPS (1 ng/ml) (N=3). Data shown are mean values ± STDEV. One-way ANOVA with Sidak’s multiple comparisons test was used in A; Two-tailed t-test was used in B and E; two-way ANOVA with Tukey’s multiple comparisons test was used in D, F and G. \**p*<0.05, \*\**p*<0.01, \*\*\**p*<0.001, \*\*\*\**p*<0.0001.

### Exogenous GM1 does not affect microglial secretion of BDNF and IGF-1, but increasing total cellular ganglioside levels results in decreased microglial secretion of BDNF

As part of their homeostatic functions, microglia produce growth factors such as BDNF and IGF-1 that provide trophic support for and regulate the activity of neurons and other brain cells [60, 61]. While both factors have been involved in neuroprotection, microglia BDNF secretion is also an established contributor to neuropathic pain [62]. We sought to determine whether endogenous ganglioside levels and exogenously administered GM1 influence the expression and secretion of these two key neurotrophic factors in primary microglia cultures. Treatment of mouse microglia with GM1 (50 µM) for 24h significantly decreased BDNF mRNA expression, but not the amount of mature BDNF secreted into the medium (Figure 6A, top and bottom panel, respectively). No changes were observed in IGF-1 mRNA expression (Figure 6A). In contrast, in microglia treated with L-*t*-PDMP (15 µM for 72 h) to increase endogenous ganglioside levels, both BDNF mRNA and protein expression, as well as the amount of this factor secreted into the medium, were significantly decreased. On the contrary, IGF-1 mRNA expression was significantly increased in these cells, without significant changes in the amount of IGF-1 protein produced (Figure 6H). In cells treated with GENZ-123346 (10 µM for 72 h) to decrease ganglioside levels, no changes were observed in BDNF mRNA or protein levels, and although IGF-1 mRNA expression was increased, IGF-1 protein levels were not (Figure 6C).

**Figure 6.**
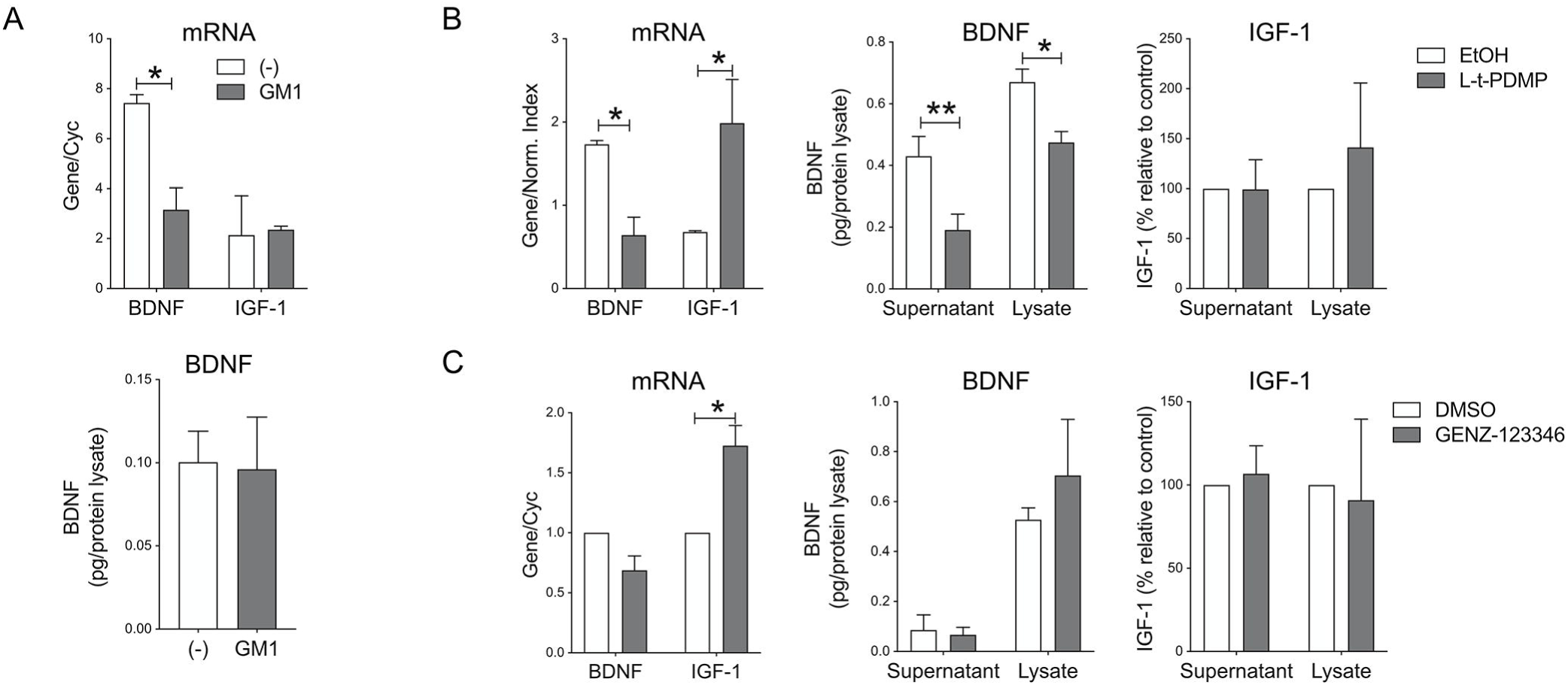
Modulation of BDNF and IGF-1 expression by gangliosides. **A)** BDNF and IGF-1 mRNA expression (top panel) and mature BDNF secreted in the medium (bottom panel) in primary murine microglia treated for 24 hours with GM1. GM1 decreases BDNF expression but not the amount of neurotrophic factor secreted into the medium. **B)** BDNF and IGF-1 expression in primary microglia treated for with L-t-PDMP (10µM for 72 h). mRNA and protein levels in total cell lysates or secreted in the medium are shown. L-t-PDMP treatment significantly decreases BDNF gene expression and protein levels, and increases IGF-1 mRNA expression (but not protein levels). **C)** BDNF and IGF-1 expression in primary microglia treated GENZ-123346 (10 µM for 72 hours). GENZ-123346 treatment does not affect BDNF expression, but increases IGF-1 mRNA expression without affecting IGF-1 protein levels. Mature BDNF and IGF-1 levels in the cell supernatant and in cell lysates were normalized to cell lysate total protein content. Graphs show mean values ± STDEV from 3 independent experiments. A-H: Two tailed unpaired *t*-test: \**p*<0.05.

### Administration of exogenous GM1 and modulation of endogenous ganglioside levels affect microglial chemotaxis and phagocytosis

Next, we sought to determine the role of gangliosides on microglial chemotaxis and phagocytosis, two microglia functions that are crucial for maintaining brain homeostasis. We performed a transwell cell migration assays using ATP as a chemotactic stimulus. As shown in Figure 7A, pre-incubation of primary microglia with 50 µM GM1 resulted in increased cell migration along the ATP gradient. This behavior was not recapitulated by treatment with L-t-PDMP (Figure 7B), suggesting that higher cellular concentrations of GM1 (as achieved through administration of exogenous ganglioside) might be required, or that the concomitant increase of other gangliosides (GD1a and GT1b, Figure 4F) upon L-t-PDMP treatment may counteract the effects of GM1 alone. On the other hand, decreasing endogenous gangliosides levels with GENZ-123346 partially inhibited microglial chemotaxis (Figure 7C). The effects of GM1 on microglial migration were further confirmed in an *in vitro* scratch assay using BV2 cells (Figure 7D) and murine primary microglia (Figure 7E). In both cases, cells treated with GM1 showed increased migratory activity. Altogether, these data indicate that chemotaxis and migratory activity of microglia is enhanced by administration of GM1, while a decrease in endogenous ganglioside levels is associated with decreased chemotaxis.

**Figure 7.**
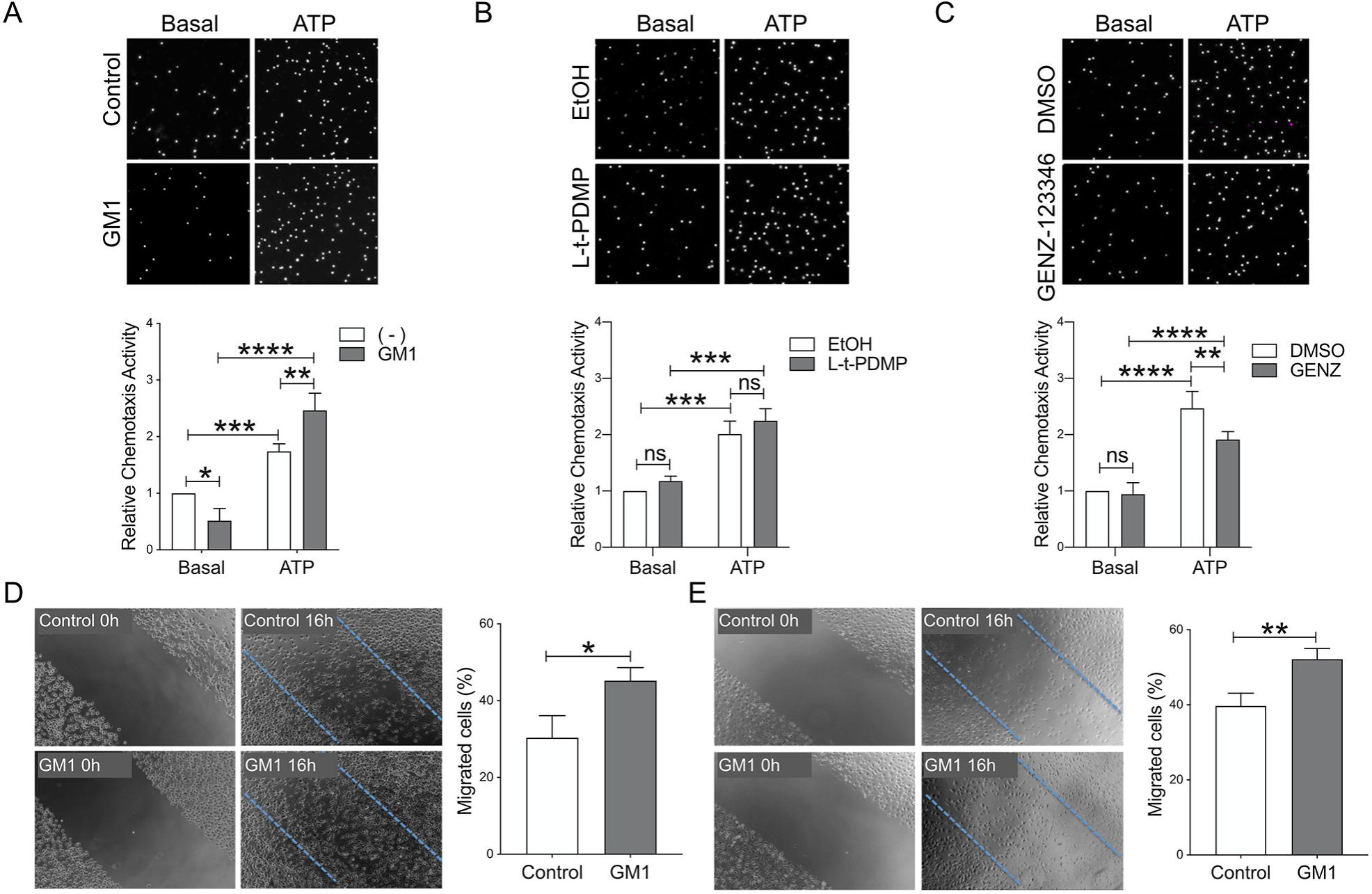
Administration of GM1 and cellular ganglioside levels affect ATP-induced chemotaxis in primary microglia. Rat primary microglia were pre-incupated with exogenous GM1 (50 μM for 4h; N=4) (**A**), L-t-PDMP (10 μM for 3 days; N=3) (**B**), or GENZ-123346 (5 μM for 3 days; N=4) (**C**). Migration towards ATP (100 ng/ml) in a transwell system and in a 2h-period was measured and normalized over cell migration in basal conditions (no ATP). Representative images of microglia migrating to the bottom side of the transwell insert and quantification of migrating microglia are shown. GM1 increased microglia chemotaxis activity and GENZ-123346 decreased it, while L-t-PDMP had no effect. **D and E)** Representative images and relative quantification of migrating BV2 cells (D) and mouse microglia (E) in a scratch assay, 16 and 24h post scratch, respectively. In both cases cells were incubated with GM1 (50 μM for 4h) prior to scratching (N=3). Data shown are mean values ± STDEV. A-C: two-way ANOVA with Tukey’s multiple comparisons test; D-E: two-tailed *t*-test; \**p*<0.05, \*\**p*<0.01, \*\*\**p*<0.001, \*\*\*\**p*<0.0001.

Microglial phagocytic activity towards latex beads was significant increased by cell pre-treatment with GM1 (50 μM for 4h), but not L-t-PDMP (10 μM for 3 days), while treatment with GENZ-123346 (10 µM for 3 days) resulted in reduced phagocytic activity (Figure 8A). In health and disease conditions, the phagocytic activity of microglia towards apoptotic bodies is crucial to maintain homeostasis and prevent overt inflammation. To determine whether ganglioside levels specifically affect the ability of microglia to clear apoptotic bodies, the latter were prepared from neuronal N2a cells and labeled with the pH-sensitive dye pHrodo Red, which enables the quantification of particles in post-phagocytic phagolysosomes. Microglia pre-treated with GM1 displayed a modest but significant increase in their ability to take-up apoptotic bodies, while modulation of cellular ganglioside levels with L-t-PDMP or GENZ-123346 had no significant effects (Figure 8B).

**Figure 8.**
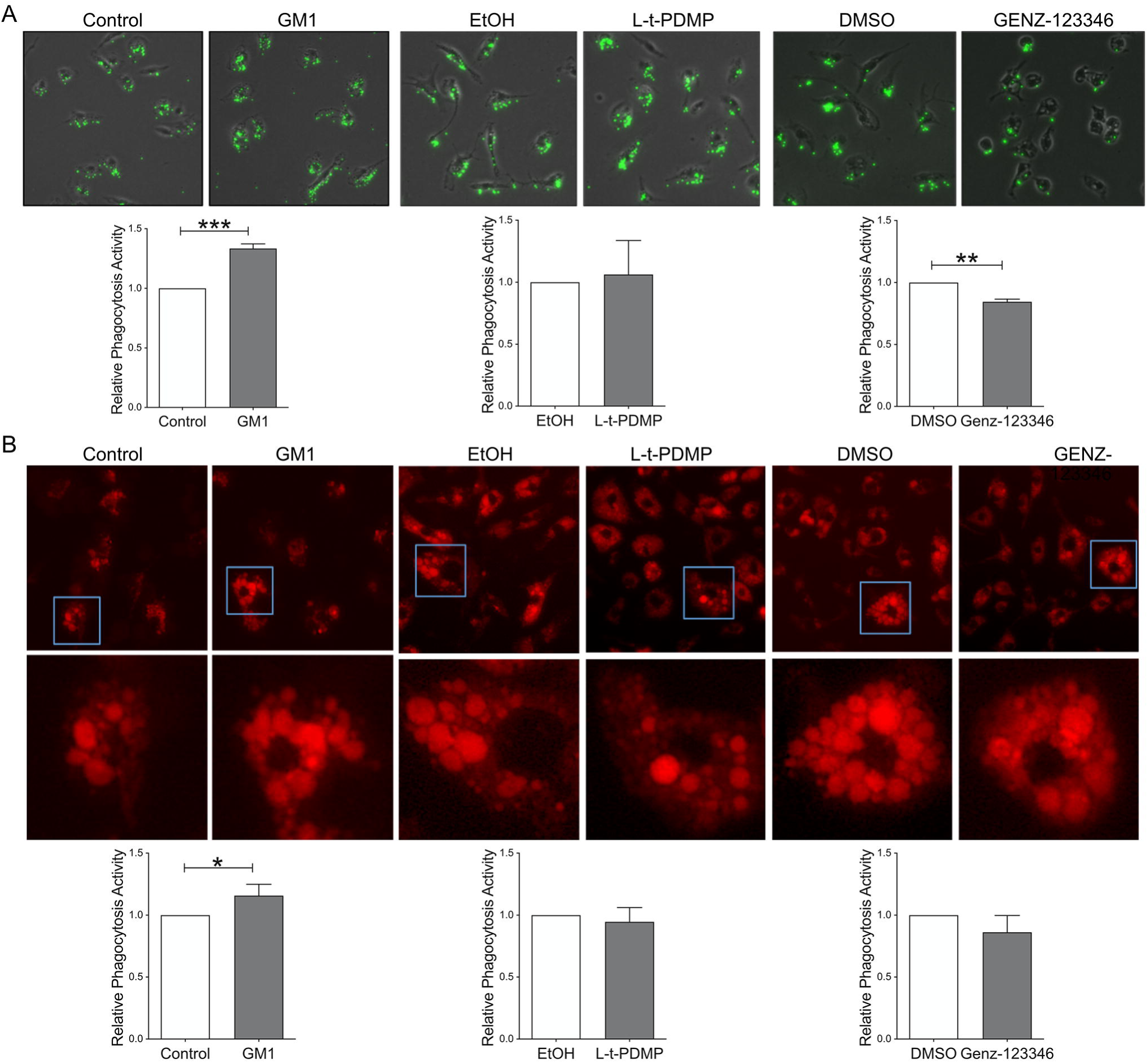
Gangliosides modulate microglia phagocytic activity towards synthetic beads and apoptotic bodies. **A)** Representative images and quantification of fluorescent latex beads uptake in rat microglia treated with GM1 (50 μM for 4h; N=4) L-t-PDMP (10 μM for 3 days; N=3) or GENZ-123346 (10 µM for 3 days; N=3). GM1 increased and GENZ-123346 decreased uptake, while treatment with L-t-PDMP had no effects. **B)** Representative images and quantification of uptake of apoptotic bodies prepared from N2a neuronal cells and labeled with pHrodo by rat microglia, following the treatments indicated in A). Bar graphs show mean values ± STDEV. Two-tailed paired *t*-test; \**p*<0.05, \*\**p*<0.01, \*\*\**p*<0.001.

### Discussion

In this study we analyzed the effects of pharmacological treatments that modulate cellular ganglioside levels on microglia functions that are crucial to healthy brain homeostasis and that represent potential targets for therapeutic intervention in disease state. Of all gangliosides, GM1 is probably the most studied and its neuroprotective properties are well-known [63]. GM1 administration was shown to provide significant benefits in a variety of neurological conditions [38], including stroke [64, 65], PD [66, 67] and HD [32, 41, 42]. Some of the underlying mechanisms were neuronal-specific [63, 68, 69], but whether GM1 could also directly target microglia was not investigated. This is an important gap in knowledge, as in all conditions listed above aberrant pro-inflammatory microglia activation and impaired homeostatic microglial functions can significantly contribute to disease pathogenesis and/or progression [9, 14, 70-73]. Our work provides evidence, for the first time, of a strong anti-inflammatory effect of exogenously administered ganglioside GM1 on BV2 microglial cells and on rat, mouse and human primary microglia exposed to various pro-inflammatory stimuli, including LPS, IL-1β and phagocytosis of latex beads, while no effects of the ganglioside were observed in unstimulated conditions. These data are in contrast with previous reports that showed activation of the p38 MAPK pathway and morphological changes suggestive of microglial activation upon microglia incubation with GM1 [74], as well as microglia secretion of pro-inflammatory TNF and NO after treatment with GT1b or with a mix of brain gangliosides containing GM1 [75-77]. While we have not tested the specific effects of other gangliosides in this study, the discrepancy between our data and other studies that observed microglia activation by GM1 [74] might be due to the specific outcomes measured (cell morphology versus production of pro-inflammatory cytokines in our studies) or to different modalities of GM1 administration, since we have observed that a micellar presentation is required for GM1 to exert its anti-inflammatory activity (data not shown). On the other hand, our findings are in line with and expand on studies performed on myeloid cells that showed that GM1 and other gangliosides attenuate the response of human monocytes, THP-1 cells and RAW 264.7 macrophages to LPS [78-81] and to amyloid-β-peptide [82, 83].

The mechanism underlying the anti-inflammatory effects of GM1 are still unknown and will require further investigation. Previous studies have shown that GM1 might bind to some LPS serotypes and potentially decrease binding to TLR4 when pre-incubated with the bacterial toxin [84, 85]. The LPS serotype used in this study (E. coli O55:B5), however, does not bind GM1 [85]. In other studies, incubation of microglia with GM1 [86] or with a mix of brain gangliosides [86] resulted in downregulation of TLR-4 expression. Although in our experiments we observed slightly decreased levels of total cellular TLR-4 in microglial BV2 cells incubated for 24h with GM1, the levels of plasma membrane receptor available for binding were not significantly changed by the ganglioside. In addition, we found that GM1 could still curtail microglia inflammatory responses *<u>after</u>* microglia had been activated, and after removal of LPS. While this does not exclude potential modulatory effects of GM1 on TLR-4 signaling, it suggests that the ganglioside activates a “shut-off” pathway that helps restoring homeostatic conditions upon exposure to an inflammatory stimulus. An anti-inflammatory mechanism independent from any potential direct effect of GM1 on TLR-4 is also suggested by our observation that GM1 decreases pro-inflammatory microglia responses triggered by IL-1β as well as phagocytosis of latex beads, which was shown to lead to activation of the NLRP3 inflammasome and downstream inflammatory response [57]. Altogether, our data suggest that administration of exogenous GM1 elicits a potent cell-autonomous anti-inflammatory response in microglia, which might contribute to the neuroprotective activity of this ganglioside in models of neurodegeneration and neuroinflammation. Future studies will address the underlying mechanism(s) and will help identifying novel strategies to lower microglia activation in the context of neuroinflammatory conditions.

L-*t*-PDMP treatment to increase microglial endogenous ganglioside levels recapitulated the effects of exogenous GM1 in BV2 cells, but less so in primary microglia, which are generally more responsive to LPS stimulation than BV2 cells [87, 88]. In primary microglia, L-*t*-PDMP elicited a much weaker anti-inflammatory response compared to the administration of exogenous GM1, decreasing expression of pro-inflammatory genes only at low LPS concentrations, but failing to attenuate microglial inflammatory responses at higher LPS concentrations. This could be due to higher levels of cellular GM1 achieved upon administration of exogenous GM1. Alternatively, the concomitant increase of other gangliosides following administration of L-*t*-PDMP, in particular GT1b, which was shown to promote microglia activation [77], might counteract the anti-inflammatory properties of GM1.

Decreasing endogenous ganglioside levels with GENZ-123346 generally resulted in opposite effects compared to GM1 administration or L-t-PDMP treatment, with increased activation of the NFkB and p38 MAPK pathways and increased transcription of TNF mRNA, although the latter was not mirrored by equivalent changes in the secretion of TNF, for reasons that remain unclear. The overall microglia response to LPS depend on several intracellular signaling cascades activated by TLR4 stimulation, among them ERK1/ERK2, JNKs and p38 [89], that might be differentially affected by changes in the relative abundance and ratios of endogenous gangliosides and differentially impact gene transcription versus protein translation and secretion. One of the best characterized roles of GM1 is its ability to stimulate neurotrophin signaling in neuronal cells, including the tropomyosin receptor kinase B (TrkB)-BDNF pathway, which has been proposed as one of the major mechanisms underlying the neuroprotective activity of this ganglioside [69]. Administration of GM1 was shown to increase expression of BDNF in neuronal SH-SY5Y cells and cortical neurons *in vitro* [90] and in the nucleus accumbens *in vivo* [91]. Furthermore, GM1 can stimulate a positive feedback loop by binding and weakly activating TrkB [92-94], which in turn increases the expression of BDNF in neurons in a CREB-dependent manner [95]. TrkB receptors [96] and BDNF [97] are also expressed in microglia, where secretion of BDNF can activate a similar autocrine positive feedback loop and prolong microglia activation under inflammatory conditions [98], but the interplay between the microglial TrkB-BDNF pathway and gangliosides is not known. Our data show for the first time that both of exogenous GM1 and L-*t*-PDMP-mediated increase in endogenous ganglioside levels result in a significant decrease in BDNF mRNA expression in microglia, and substantially decreased levels of BDNF secreted by cells treated with L-*t*-PDMP. This is in stark contrast with the effects of GM1 on neuronal neurotrophin secretion, and suggests the presence of different mechanisms activated by gangliosides that modulate BDNF transcription and secretion in a cell context-dependent manner. Our data also point at a potential beneficial role of gangliosides and/or L-*t*-PDMP in neuropathic pain, where a pathological increase in the secretion of microglial BDNF results in abnormal neuronal network excitability and altered sensitivity to pain [61].

We also investigated the effects of gangliosides on the production of microglial IGF-1, a growth factor produced by both neurons and glial cells with neuroprotective, anti-inflammatory and reparative roles [5, 60, 99]. Microglia is a major source of IGF-1 in the brain [100], especially in conditions that stimulate the acquisition by microglia of an M2-like immunoregulatory/reparative phenotype [101]. We found that expression of IGF-1 mRNA was not affected by exogenous GM1. Both L-*t*-PDMP and GENZ-123346 increased mRNA, but not protein levels of IGF-1, suggesting that gangliosides do not significantly affect the ability of microglia to produce this growth factor, at least in basal conditions.

Finally, our studies show that gangliosides can significantly modulate chemotaxis and microglial phagocytic activity, two microglia functions that are crucial to healthy brain homeostasis and damage repair at sites of injury [102-104]. Treatment of microglia with GM1 significantly increased microglia chemotaxis towards ATP, a major chemotactic stimulus in injury and disease conditions [103], as well as microglia migration in a wound scratch assay. Decreasing endogenous ganglioside levels with GENZ had the opposite effect. These findings are in line with studies that showed increased cell migration of a glioma cell lines exposed to GM1 [105] and inhibition of CXCR4-dependent cell migration in a neuroepithelioma cell line upon inhibition of UGCG [106], and suggest an important role of GM1 in the response to of chemotactic stimuli. L-*t*-PDMP was not able to recapitulate the effects of exogenous GM1 on microglia chemotaxis, which suggests that the relative ratio between GM1 and other gangliosides co-increased by cell treatment with L-*t*-PDMP might be an important determinant of the overall effects on chemotaxis. In this regard, it is important to mention that GM3, a minor brain ganglioside species that was not measured in our studies and that could also increase upon cell treatment with L-*t*-PDMP, was shown to inhibit cell migration by keratinocytes [107].

Administration of exogenous GM1, but not L-t-PDMP treatment, significantly increased phagocytosis of both latex beads and apoptotic bodies by primary microglia. On the other hand, decreasing endogenous microglial ganglioside levels resulted in reduced uptake of apoptotic bodies but not latex beads, suggesting a differential modulatory action of gangliosides on different phagocytic pathways. How GM1 increases chemotaxis and phagocytosis - which are mediated by various receptors, several of which could be involved in both processes [108] - remains to be investigated. We speculate that upon insertion into cellular membranes, GM1 might promote receptor activation and/or the formation of protein complexes required for cell motility and phagocytosis [109-112].

In conclusion, our data suggest that microglial gangliosides play an important role in the regulation of crucial microglia activities - including response to inflammatory stimuli, chemotaxis and phagocytosis - that might be compromised as a result of decreased ganglioside synthesis in neurodegenerative disorders such as PD [34, 113] and HD [32, 33]. Our studies provide important insights into the therapeutic role of GM1 in neurodegenerative disease and demonstrate that the ganglioside can target inflammatory microglia in addition to neurons. These findings expand the potential of GM1-based therapies to the treatment of neuroinflammatory diseases.

## ACKNOWLEDGEMENTS

This work was supported by grants from the Natural Sciences and Engineering Research Council of Canada (NSERC), from Brain Canada/Huntington Society of Canada and from the Alberta Glycomics Centre. Experiments were performed at the University of Alberta Faculty of Medicine and Dentistry Flow Cytometry Core, which receives financial support from the Faculty of Medicine and Dentistry and Canada Foundation for Innovation (CF) awards to contributing investigators.

## REFERENCES

1. Frost, J.L. and D.P. Schafer, Microglia: Architects of the Developing Nervous System. Trends Cell Biol, 2016. 26(8): p. 587–597.

2. Li, Q. and B.A. Barres, Microglia and macrophages in brain homeostasis and disease. Nat Rev Immunol, 2018. 18(4): p. 225–242.

3. Paolicelli, R.C., et al., Synaptic pruning by microglia is necessary for normal brain development. Science, 2011. 333(6048): p. 1456–8.

4. Schafer, D.P., et al., Microglia sculpt postnatal neural circuits in an activity and complement-dependent manner. Neuron, 2012. 74(4): p. 691–705.

5. Ueno, M., et al., Layer V cortical neurons require microglial support for survival during postnatal development. Nat Neurosci, 2013. 16(5): p. 543–51.

6. Parkhurst, C.N., et al., Microglia promote learning-dependent synapse formation through brain-derived neurotrophic factor. Cell, 2013. 155(7): p. 1596–609.

7. Cunningham, C.L., V. Martinez-Cerdeno, and S.C. Noctor, Microglia regulate the number of neural precursor cells in the developing cerebral cortex. J Neurosci, 2013. 33(10): p. 4216–33.

8. Luo, X.G., J.Q. Ding, and S.D. Chen, Microglia in the aging brain: relevance to neurodegeneration. Mol Neurodegener, 2010. 2010: p. 12.

9. Lucin, K.M. and T. Wyss-Coray, Immune activation in brain aging and neurodegeneration: too much or too little? Neuron, 2009. 64(1): p. 110–22.

10. Heneka, M.T., M.P. Kummer, and E. Latz, Innate immune activation in neurodegenerative disease. Nat Rev Immunol, 2014. 14(7): p. 463–77.

11. Ransohoff, R.M., How neuroinflammation contributes to neurodegeneration. Science, 2016. 353(6301): p. 777–83.

12. Heneka, M.T., et al., Neuroinflammation in Alzheimer’s disease. Lancet Neurol, 2015. 14(4): p. 388–405.

13. Crotti, A. and C.K. Glass, The choreography of neuroinflammation in Huntington’s disease. Trends Immunol, 2015. 36(6): p. 364–73.

14. Bjorkqvist, M., et al., A novel pathogenic pathway of immune activation detectable before clinical onset in Huntington’s disease. J Exp Med, 2008. 205(8): p. 1869–77.

15. McGeer, P.L. and E.G. McGeer, Glial reactions in Parkinson’s disease. Mov Disord, 2008. 23(4): p. 474–83.

16. More, S.V., et al., Cellular and molecular mediators of neuroinflammation in the pathogenesis of Parkinson’s disease. Mediators Inflamm, 2013. 2013: p. 952375.

17. Zhou, J.Y., et al., The Glycoscience of Immunity. Trends Immunol, 2018. 39(7): p. 523–535.

18. Fukuda, M., U. Rutishauser, and R.L. Schnaar, Neuroglycobiology. Molecular and cellular neurobiology series. 2005, Oxford ; New York: Oxford University Press. xix, 229 p.

19. Schnaar, R.L. and T. Kinoshita, Glycosphingolipids, in Essentials of Glycobiology, rd, et al., Editors. 2015, Cold Spring Harbor Laboratory Press Copyright 2015-2017 by The Consortium of Glycobiology Editors, La Jolla, California. All rights reserved.: Cold Spring Harbor (NY). p. 125–135.

20. Posse de Chaves, E. and S. Sipione, Sphingolipids and gangliosides of the nervous system in membrane function and dysfunction. FEBS Lett, 2010. 584(9): p. 1748–59.

21. Simpson, M.A., et al., Infantile-onset symptomatic epilepsy syndrome caused by a homozygous loss-of-function mutation of GM3 synthase. Nat Genet, 2004. 36(11): p. 1225–9.

22. Fragaki, K., et al., Refractory epilepsy and mitochondrial dysfunction due to GM3 synthase deficiency. Eur J Hum Genet, 2013. 21(5): p. 528–34.

23. Boccuto, L., et al., A mutation in a ganglioside biosynthetic enzyme, ST3GAL5, results in salt & pepper syndrome, a neurocutaneous disorder with altered glycolipid and glycoprotein glycosylation. Hum Mol Genet, 2014. 23(2): p. 418–33.

24. Wakil, S.M., et al., Novel B4GALNT1 mutations in a complicated form of hereditary spastic paraplegia. Clin Genet, 2014. 86(5): p. 500–1.

25. Harlalka, G.V., et al., Mutations in B4GALNT1 (GM2 synthase) underlie a new disorder of ganglioside biosynthesis. Brain, 2013. 136(Pt 12): p. 3618–24.

26. Boukhris, A., et al., Alteration of ganglioside biosynthesis responsible for complex hereditary spastic paraplegia. Am J Hum Genet, 2013. 93(1): p. 118–23.

27. Allende, M.L. and R.L. Proia, Simplifying complexity: genetically resculpting glycosphingolipid synthesis pathways in mice to reveal function. Glycoconj J, 2014. 31(9): p. 613–22.

28. Kracun, I., et al., Gangliosides in the human brain development and aging. Neurochem Int, 1992. 20(3): p. 421–31.

29. Mo, L., et al., GM1 and ERK signaling in the aged brain. Brain Res, 2005. 1054(2): p. 125–34.

30. Palestini, P., et al., Changes in the ceramide composition of rat forebrain gangliosides with age. J Neurochem, 1990. 54(1): p. 230–5.

31. Segler-Stahl, K., J.C. Webster, and E.G. Brunngraber, Changes in the concentration and composition of human brain gangliosides with aging. Gerontology, 1983. 29(3): p. 161–8.

32. Maglione, V., et al., Impaired ganglioside metabolism in Huntington’s disease and neuroprotective role of GM1. J Neurosci, 2010. 30(11): p. 4072–80.

33. Desplats, P.A., et al., Glycolipid and ganglioside metabolism imbalances in Huntington’s disease. Neurobiol Dis, 2007. 27(3): p. 265–77.

34. Wu, G., et al., Deficiency of ganglioside GM1 correlates with Parkinson’s disease in mice and humans. J Neurosci Res, 2012. 90(10): p. 1997–2008.

35. Blennow, K., et al., Gangliosides in cerebrospinal fluid in ‘probable Alzheimer’s disease’. Arch Neurol, 1991. 48(10): p. 1032–5.

36. Blennow, K., et al., Differences in cerebrospinal fluid gangliosides between “probable Alzheimer’s disease” and normal aging. Aging (Milano), 1992. 4(4): p. 301–6.

37. Alpaugh M, G.D., Forero J, Morales LC, Lackey S, Kar P, Di Pardo A, Holt A, Kerr B, Todd K, Baker GB, Fouad K and Sipione S., Therapeutic and disease-modifying effects of ganglioside GM1 in mouse models of Huntington’s disease., in Submitted. 2017.

38. Magistretti, P.J., et al., Gangliosides: Treatment Avenues in Neurodegenerative Disease. Front Neurol, 2019. 2019: p. 859.

39. Pope-Coleman, A. and J.S. Schneider, Effects of Chronic GM1 Ganglioside Treatment on Cognitieve and Motor Deficits in a Slowly Progressing Model of Parkinsonism in Non-Human Primates. Restor Neurol Neurosci, 1998. 12(4): p. 255–266.

40. Schneider, J.S., GM1 ganglioside in the treatment of Parkinson’s disease. Ann N Y Acad Sci, 1998. 1998: p. 363–73.

41. Alpaugh, M., et al., Disease-modifying effects of ganglioside GM1 in Huntington’s disease models. EMBO Mol Med, 2017. 9(11): p. 1537–1557.

42. Di Pardo, A., et al., Ganglioside GM1 induces phosphorylation of mutant huntingtin and restores normal motor behavior in Huntington disease mice. Proc Natl Acad Sci U S A, 2012. 109(9): p. 3528–33.

43. Ohmi, Y., et al., Gangliosides are essential in the protection of inflammation and neurodegeneration via maintenance of lipid rafts: elucidation by a series of ganglioside-deficient mutant mice. J Neurochem, 2011. 116(5): p. 926–35.

44. Ohmi, Y., et al., Ganglioside deficiency causes inflammation and neurodegeneration via the activation of complement system in the spinal cord. J Neuroinflammation, 2014. 2014: p. 61.

45. Gong, G., et al., Ganglioside GM1 protects against high altitude cerebral edema in rats by suppressing the oxidative stress and inflammatory response via the PI3K/AKT-Nrf2 pathway. Mol Immunol, 2018. 2018: p. 91–98.

46. Saura, J., J.M. Tusell, and J. Serratosa, High-yield isolation of murine microglia by mild trypsinization. Glia, 2003. 44(3): p. 183–9.

47. Rabinovitch, M. and M.J. DeStefano, Use of the local anesthetic lidocaine for cell harvesting and subcultivation. In Vitro, 1975. 11(6): p. 379–81.

48. Walsh, J.G., et al., Rapid inflammasome activation in microglia contributes to brain disease in HIV/AIDS. Retrovirology, 2014. 2014: p. 35.

49. Blasi, E., et al., Immortalization of murine microglial cells by a v-raf/v-myc carrying retrovirus. J Neuroimmunol, 1990. 27(2-3): p. 229–37.

50. Vandesompele, J., et al., Accurate normalization of real-time quantitative RT-PCR data by geometric averaging of multiple internal control genes. Genome Biol, 2002. 3(7): p. RESEARCH0034.

51. Liang, C.C., A.Y. Park, and J.L. Guan, In vitro scratch assay: a convenient and inexpensive method for analysis of cell migration in vitro. Nat Protoc, 2007. 2(2): p. 329–33.

52. Franklin, T.C., C. Xu, and R.S. Duman, Depression and sterile inflammation: Essential role of danger associated molecular patterns. Brain Behav Immun, 2018. 2018: p. 2–13.

53. Park, J.S., et al., High mobility group box 1 protein interacts with multiple Toll-like receptors. Am J Physiol Cell Physiol, 2006. 290(3): p. C917–24.

54. Azam, S., et al., Regulation of Toll-Like Receptor (TLR) Signaling Pathway by Polyphenols in the Treatment of Age-Linked Neurodegenerative Diseases: Focus on TLR4 Signaling. Front Immunol, 2019. 2019: p. 1000.

55. Glass, C.K., et al., Mechanisms underlying inflammation in neurodegeneration. Cell, 2010. 140(6): p. 918–34.

56. Voet, S., et al., Inflammasomes in neuroinflammatory and neurodegenerative diseases. EMBO Mol Med, 2019. 11(6).

57. Rajamaki, K., et al., Cholesterol crystals activate the NLRP3 inflammasome in human macrophages: a novel link between cholesterol metabolism and inflammation. PLoS One, 2010. 5(7): p. e11765.

58. Inokuchi, J., S. Usuki, and M. Jimbo, Stimulation of glycosphingolipid biosynthesis by L-threo-1-phenyl-2-decanoylamino-1-propanol and its homologs in B16 melanoma cells. J Biochem, 1995. 117(4): p. 766–73.

59. Shayman, J.A., The design and clinical development of inhibitors of glycosphingolipid synthesis: will invention be the mother of necessity? Trans Am Clin Climatol Assoc, 2013. 2013: p. 46–60.

60. Labandeira-Garcia, J.L., et al., Insulin-Like Growth Factor-1 and Neuroinflammation. Front Aging Neurosci, 2017. 2017: p. 365.

61. Ferrini, F. and Y. De Koninck, Microglia control neuronal network excitability via BDNF signalling. Neural Plast, 2013. 2013: p. 429815.

62. Trang, T., S. Beggs, and M.W. Salter, Brain-derived neurotrophic factor from microglia: a molecular substrate for neuropathic pain. Neuron Glia Biol, 2011. 7(1): p. 99–108.

63. Ledeen, R.W. and G. Wu, The multi-tasked life of GM1 ganglioside, a true factotum of nature. Trends Biochem Sci, 2015. 40(7): p. 407–18.

64. Oppenheimer, S., GM1 ganglioside therapy in acute ischemic stroke. Stroke, 1990. 21(5): p. 825.

65. Simon, R.P., J. Chen, and S.H. Graham, GM1 ganglioside treatment of focal ischemia: a dose-response and microdialysis study. J Pharmacol Exp Ther, 1993. 265(1): p. 24–9.

66. Pope-Coleman, A., J.P. Tinker, and J.S. Schneider, Effects of GM1 ganglioside treatment on pre- and postsynaptic dopaminergic markers in the striatum of parkinsonian monkeys. Synapse, 2000. 36(2): p. 120–8.

67. Schneider, J.S., et al., A randomized, controlled, delayed start trial of GM1 ganglioside in treated Parkinson’s disease patients. J Neurol Sci, 2013. 324(1-2): p. 0140–8.

68. Ledeen, R.W. and G. Wu, Gangliosides, alpha-Synuclein, and Parkinson’s Disease. Prog Mol Biol Transl Sci, 2018. 2018: p. 435–454.

69. Mocchetti, I., Exogenous gangliosides, neuronal plasticity and repair, and the neurotrophins. Cell Mol Life Sci, 2005. 62 (19-20): p. 2283–94.

70. Perry, V.H., J.A. Nicoll, and C. Holmes, Microglia in neurodegenerative disease. Nat Rev Neurol, 2010. 6(4): p. 193–201.

71. Kwan, W., et al., Mutant huntingtin impairs immune cell migration in Huntington disease. J Clin Invest, 2012. 122(12): p. 4737–47.

72. Crotti, A., et al., Mutant Huntingtin promotes autonomous microglia activation via myeloid lineage-determining factors. Nat Neurosci, 2014. 17(4): p. 513–21.

73. Moller, T., Neuroinflammation in Huntington’s disease. J Neural Transm, 2010. 117(8): p. 1001–8.

74. Park, J.Y., et al., GM1 induces p38 and microtubule dependent ramification of rat primary microglia in vitro. Brain Res, 2008. 2008: p. 13–23.

75. Min, K.J., et al., Gangliosides activate microglia via protein kinase C and NADPH oxidase. Glia, 2004. 48(3): p. 197–206.

76. Min, K.J., et al., Protein kinase A mediates microglial activation induced by plasminogen and gangliosides. Exp Mol Med, 2004. 36(5): p. 461–7.

77. Pyo, H., et al., Gangliosides activate cultured rat brain microglia. J Biol Chem, 1999. 274(49): p. 34584–9.

78. Ziegler-Heitbrock, H.W., et al., Gangliosides suppress tumor necrosis factor production in human monocytes. J Immunol, 1992. 148(6): p. 1753–8.

79. Wang, Y., et al., Ganglioside GD1a suppresses LPS-induced pro-inflammatory cytokines in RAW264.7 macrophages by reducing MAPKs and NF-kappaB signaling pathways through TLR4. Int Immunopharmacol, 2015. 28(1): p. 136–45.

80. Shen, W., et al., Inhibition of TLR Activation and Up-Regulation of IL-1R-Associated Kinase-M Expression by Exogenous Gangliosides. The Journal of Immunology, 2008. 180(7): p. 4425--4432.

81. Wang, Y., et al., Ganglioside GD1a suppresses LPS-induced pro-inflammatory cytokines in RAW264.7 macrophages by reducing MAPKs and NF-$\kappa$B signaling pathways through TLR4. International Immunopharmacology, 2015. 28(1): p. 136--145.

82. Ariga, T. and R.K. Yu, GM1 inhibits amyloid beta-protein-induced cytokine release. Neurochem Res, 1999. 24(2): p. 219–26.

83. Ariga, T., et al., Gangliosides inhibit the release of interleukin-1beta in amyloid beta-protein-treated human monocytic cells. J Mol Neurosci, 2001. 17(3): p. 371–7.

84. Cavaillon, J.M., et al., Inhibition by gangliosides of the specific binding of lipopolysaccharide (LPS) to human monocytes prevents LPS-induced interleukin-1 production. Cell Immunol, 1987. 106(2): p. 293–303.

85. Jeng, K.C., T.L. Chen, and J.L. Lan, Gangliosides suppression of murine lymphoproliferation and interleukin 1 production. Immunol Lett, 1988. 19(4): p. 335–40.

86. Jou, I., et al., Gangliosides trigger inflammatory responses via TLR4 in brain glia. Am J Pathol, 2006. 168(5): p. 1619–30.

87. Das, A., et al., Transcriptome sequencing reveals that LPS-triggered transcriptional responses in established microglia BV2 cell lines are poorly representative of primary microglia. Journal of Neuroinflammation, 2016. 13(1): p. 182.

88. Henn, A., et al., The suitability of BV2 cells as alternative model system for primary microglia cultures or for animal experiments examining brain inflammation. Altex, 2009. 26(2): p. 83–94.

89. van der Bruggen, T., et al., Lipopolysaccharide-induced tumor necrosis factor alpha production by human monocytes involves the raf-1/MEK1-MEK2/ERK1-ERK2 pathway. Infect Immun, 1999. 67(8): p. 3824–9.

90. Shin, M.K., et al., The ganglioside GQ1b regulates BDNF expression via the NMDA receptor signaling pathway. Neuropharmacology, 2014. 2014: p. 414–21.

91. Valdomero, A., et al., Exogenous GM1 ganglioside increases accumbal BDNF levels in rats. Behav Brain Res, 2015. 2015: p. 303–6.

92. Pitto, M., et al., Influence of endogenous GM1 ganglioside on TrkB activity, in cultured neurons. FEBS Lett, 1998. 439(1-2): p. 93–6.

93. Bachis, A., et al., Gangliosides prevent excitotoxicity through activation of TrkB receptor. Neurotox Res, 2002. 4(3): p. 225–34.

94. Duchemin, A.M., et al., GM1 ganglioside induces phosphorylation and activation of Trk and Erk in brain. J Neurochem, 2002. 81(4): p. 696–707.

95. Esvald, E.E., et al., CREB Family Transcription Factors Are Major Mediators of BDNF Transcriptional Autoregulation in Cortical Neurons. J Neurosci, 2020. 40(7): p. 1405–1426.

96. Condorelli, D.F., et al., Neurotrophins and their trk receptors in cultured cells of the glial lineage and in white matter of the central nervous system. J Mol Neurosci, 1995. 6(4): p. 237–48.

97. Gomes, C., et al., Activation of microglial cells triggers a release of brain-derived neurotrophic factor (BDNF) inducing their proliferation in an adenosine A2A receptor-dependent manner: A2A receptor blockade prevents BDNF release and proliferation of microglia. Journal of Neuroinflammation, 2013. 10(1): p. 780.

98. Zhang, X., et al., Positive Feedback Loop of Autocrine BDNF from Microglia Causes Prolonged Microglia Activation. Cellular Physiology and Biochemistry, 2014. 34(3): p. 715–723.

99. Bellver-Landete, V., et al., Microglia are an essential component of the neuroprotective scar that forms after spinal cord injury. Nat Commun, 2019. 10(1): p. 518.

100. Rodriguez-Perez, A.I., et al., Crosstalk between insulin-like growth factor-1 and angiotensin-II in dopaminergic neurons and glial cells: role in neuroinflammation and aging. Oncotarget, 2016. 7(21): p. 30049–67.

101. Ferger, A.I., et al., Effects of mitochondrial dysfunction on the immunological properties of microglia. J Neuroinflammation, 2010. 2010: p. 45.

102. Nimmerjahn, A., F. Kirchhoff, and F. Helmchen, Resting microglial cells are highly dynamic surveillants of brain parenchyma in vivo. Science, 2005. 308(5726): p. 1314–8.

103. Davalos, D., et al., ATP mediates rapid microglial response to local brain injury in vivo. Nat Neurosci, 2005. 8(6): p. 752–8.

104. Galloway, D.A., et al., Phagocytosis in the Brain: Homeostasis and Disease. Front Immunol, 2019. 2019: p. 790.

105. Merzak, A., et al., Gangliosides modulate proliferation, migration, and invasiveness of human brain tumor cells in vitro. Mol Chem Neuropathol, 1995. 24(2-3): p. 121–35.

106. Limatola, C., et al., Evidence for a role of glycosphingolipids in CXCR4-dependent cell migration. FEBS Lett, 2007. 581(14): p. 2641–6.

107. Dam, D.H.M., et al., Ganglioside GM3 Mediates Glucose-Induced Suppression of IGF-1 Receptor-Rac1 Activation to Inhibit Keratinocyte Motility. J Invest Dermatol, 2017. 137(2): p. 440–448.

108. Noda, M. and A. Suzumura, Sweepers in the CNS: Microglial Migration and Phagocytosis in the Alzheimer Disease Pathogenesis. Int J Alzheimers Dis, 2012. 2012: p. 891087.

109. Jawhara, S., et al., Distinct Effects of Integrins αXβ2 and αMβ2 on Leukocyte Subpopulations during Inflammation and Antimicrobial Responses. Infection and immunity, 2016. 85(1): p. e00644–16.

110. Rosales, C. and E. Uribe-Querol, Phagocytosis: A Fundamental Process in Immunity. BioMed research international, 2017. 2017: p. 9042851–9042851.

111. Gómez-Moutón, C.n., et al., Dynamic redistribution of raft domains as an organizing platform for signaling during cell chemotaxis. Journal of Cell Biology, 2004. 164(5): p. 759–768.

112. Gómez-Moutón, C., et al., Segregation of leading-edge and uropod components into specific lipid rafts during T cell polarization. Proceedings of the National Academy of Sciences, 2001. 98(17): p. 9642–9647.

113. Schneider, J.S., Altered expression of genes involved in ganglioside biosynthesis in substantia nigra neurons in Parkinson’s disease. PLoS One, 2018. 13(6): p. e0199189.

