## Supplemental materials and figures for "Anti-inflammatory role of GM1 and modulatory effects of gangliosides on microglia functions"

#### Supplementary Methods

**Analysis of cell viability and TLR-4 surface expression by flow cytometry.**

Cells viability of primary microglia cells was measured using Annexin V-PE (BD Pharmigen 556421) and LIVE/DEAD Fixable Far Red Dead Cell Stain Kit (Invitrogen L34974). Briefly, cells were detached from the plates using TryPLE Express Enzyme (Gibco), washed with cold PBS and stained for 15 minutes with Anexin V-PE and LIVE/DEAD Fixable Far Red. After washing in PBS, fluorescence emission was detected using an Attune NxT Flow Cytometer (Invitrogen). To quantify TLR4 expression at the plasma membrane, BV-2 cells were incubated with 1% BSA on ice for 30 minutes and then stained with mouse anti-TLR4 (Abcam, ab22048) without fixation nor permeabilization. After washing, Alexa Fluor-488 anti-mouse IgG (Invitrogen) was added for 1h on ice. Unstained controls were prepared by incubation with secondary antibodies only. Cells were analyzed with BD FACS Canto II (Becton Dickinson). Flow cytometry data were analyzed using FlowJo software.

#### Supplementary Figure Legends

**Supplementary Figure 1**

**Expression of TLR-4 in BV-2 cells after GM1, L-t-PDMP or GENZ-123346 administration.** **A)** Representative Western blot and quantification total TLR4 expression in BV-2 cells treated with GM1 (50 µM) for 4 and 24 h. Data are mean values ± STDEV from 4 independent experiments. One-way ANOVA with Dunnett’s multiple comparisons test; **p*<0.05. B) Representative histogram and flow cytometry quantification (mean fluorescence intensity, MFI) of TLR4 present at the plasma membrane of BV-2 cells after treatment with GM1 (as indicated in A), 10 µM L-*t*-PDMP or 10µM GENZ-123346 for 72 hours. All histograms stain

**Supplementary Figure 2**

**Treatment with exogenous GM1 does not affect microglia survival. A)** Cell viability assay using LIVE/DEAD™ Fixable Near-IR Dead Cells stain and Annexin-V PE in mouse microglia pre-incubated with GM1 (50 µM for 1 h) and then stimulated with LPS (100 ng/ml for 24 h), in the presence or absence of GM1. **B)** Cell viability assay in mouse microglia stimulated with LPS (100 ng/ml) for 3 h, washed and then incubated with GM1 (50μM) for 8 additional hours.

### Figure S1

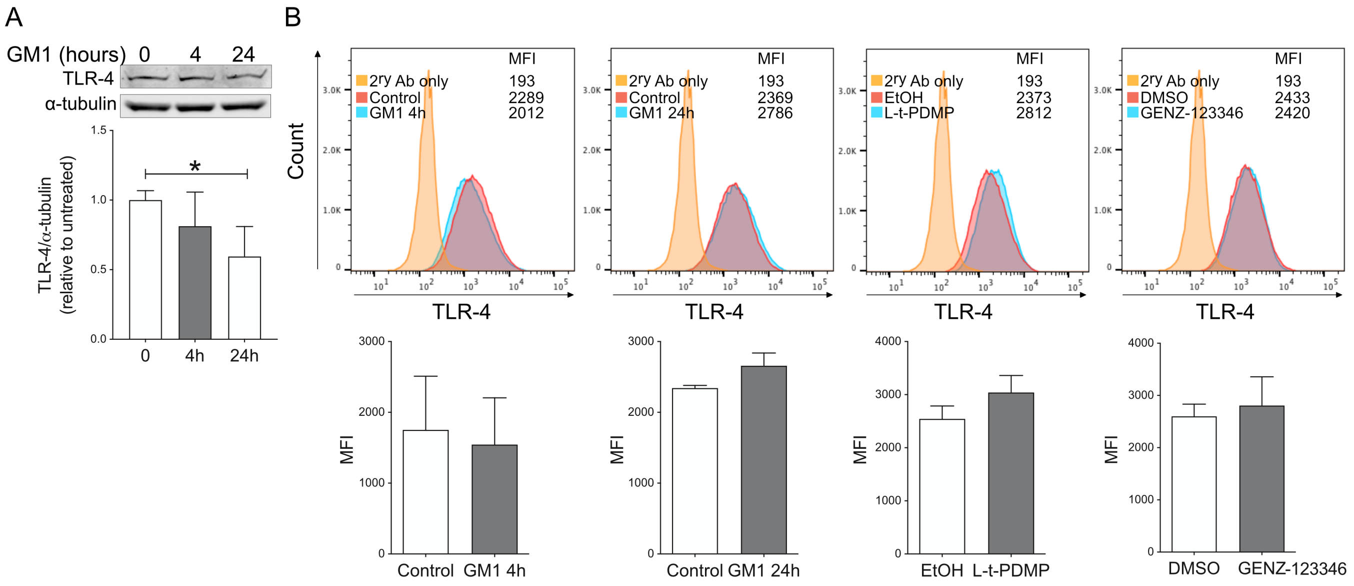

A
GM1 (hours) 0 4 24
TLR-4
α-tubulin
TLR-4/α-tubulin (relative to untreated)
1.5
1.0
0.5
0.0
*
0
4h
24h
B
Count
MFI
2ry Ab only 193
Control 2289
GM1 4h 2012
2ry Ab only 193
Control 2369
GM1 24h 2786
2ry Ab only 193
EtOH 2373
L-t-PDMP 2812
2ry Ab only 193
DMSO 2433
GENZ-123346 2420
TLR-4
MFI
Control
GM1 4h
Control
GM1 24h
EtOH
L-t-PDMP
DMSO
GENZ-123346

### Figure S2

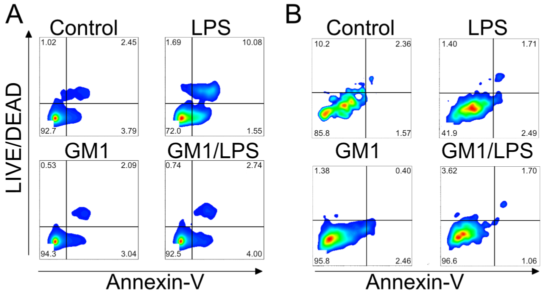
A
Control
LPS
GM1
GM1/LPS
LIVE/DEAD
Annexin-V
B
Control
LPS
GM1
GM1/LPS
Annexin-V
